# *Salmonella* Typhimurium infection inhibits macrophage IFNβ signaling in a TLR4-dependent manner

**DOI:** 10.1101/2024.03.05.583530

**Authors:** Michael Shuster, Zhihui Lyu, Jacques Augenstreich, Shrestha Mathur, Akshaya Ganesh, Jiqiang Ling, Volker Briken

## Abstract

Type I Interferons (IFNs) generally have a protective role during viral infections, but their function during bacterial infections is dependent on the bacterial species. *Legionella pneumophila*, *Shigella sonnei* and *Mycobacterium tuberculosis* can inhibit type I IFN signaling. Here we examined the role of type I IFN, specifically IFNβ, in the context of *Salmonella enterica* serovar Typhimurium (STm) macrophage infections and the capacity of STm to inhibit type I IFN signaling. We demonstrate that IFNβ has no effect on the intracellular growth of STm in infected bone marrow derived macrophages (BMDMs) derived from C57BL/6 mice. STm infection inhibits IFNβ signaling but not IFNγ signaling in a murine macrophage cell line. We show that this inhibition is independent of the type III and type VI secretion systems expressed by STm and is also independent of bacterial phagocytosis. The inhibition is Toll-like receptor 4 (TLR4)-dependent as the TLR4 ligand, lipopolysaccharide (LPS), alone is sufficient to inhibit IFNβ-mediated signaling and STm-infected, TLR4-deficient BMDMs do not exhibit inhibited IFNβ signaling. In summary, we show that macrophages exposed to STm have reduced IFNβ signaling via crosstalk with TLR4 signaling, and that IFNβ signaling does not affect cell autonomous host defense against STm.

## Introduction

*Salmonella enterica* is a Gram-negative, facultative intracellular pathogen that causes enteric infections leading to gastroenteritis, typically caused by nontyphoidal serovars, or to typhoid and paratyphoid fevers, caused by typhoidal serovars^1^. *Salmonella* makes extensive use of type III secretion systems (T3SS) to establish infection and manipulate host cells. These secretion systems are encoded in *Salmonella* pathogenicity islands (SPIs). The SPI-1 T3SS injects effectors across host cell plasma membranes to facilitate invasion of epithelial cells and trigger inflammation^2–6^. Once inside the host cell, effectors secreted by the SPI-2 T3SS establish a replicative niche by maintaining vacuole stability and disrupting host cell functions such as endolysosomal trafficking and inflammatory cell death^7^. Type VI secretion systems (T6SSs) transport effectors into neighboring bacterial and eukaryotic cells^8–10^. The T6SS encoded on SPI-6 is involved in gut colonization^11^, interbacterial competition^12^, and intracellular replication^13^ of *Salmonella*.

Type I interferons (IFNs) are comprised of IFNα (13 isoforms), IFNβ, as well as several less characterized ones such as IFNδ and IFNκ. They signal through a unique heterodimeric IFNα/β receptor (IFNAR). Trimerization of type I IFN and receptor subunits induces signal activation via tyrosine phosphorylation of Janus kinases, JAK1 and TYK2, which are associated with the intracellular domains of IFNAR^14,15^. Upon activation, JAK1 and TYK2 phosphorylate STAT1 and STAT2. These phosphorylated STAT proteins, upon binding to IRF9, assemble to form the trimeric transcription factor known as ISGF3. ISGF3 translocates to the nucleus and binds to a consensus interferon stimulated response element (ISRE) in the promotor region of IFN-stimulated genes (ISGs)^16,17^. This induces the transcription of hundreds of genes that regulate different aspects of host immunity^18–20^. Type I IFN signaling is a tightly regulated process that involves the aforementioned activation pathway as well as negative regulators such as suppressor of cytokine signaling (SOCS) proteins^21,22^ and ubiquitin-specific peptidase 18 (USP18)^22–24^

Type I IFNs are largely known for their essential role in antiviral immunity^25–27^. The selective pressure on viruses led to the evolution of many viral proteins with the ability to inhibit host cell type I IFN signaling^28–30^. In contrast, the impact of type I IFNs on host responses to bacterial pathogens varies, exhibiting both beneficial and detrimental effects that is dependent on the bacterial species and the experimental model (e.g., cell type or animal) employed in the study^31–33^. For example, in *Mycobacterium tuberculosis* infections, type I IFN proves detrimental^34,35^, while it is linked to host protection in Legionella pneumophila infections^31–33^. In the context of *Salmonella* infections, type I IFN displays both beneficial^36–38^ and detrimental^39–44^ effects. In the context of *ex vivo* epithelial cell infections, type I IFN is protective by reinforcing the epithelial barrier^38^. Similar to many viruses, some bacterial pathogens such as *M. tuberculosis*^45–47^, *Shigella sonnei*^48^, and *L. pneumophila*^49^ inhibit type I IFN signaling. *M. tuberculosis* specifically inhibits type I IFN signaling at the level of JAK1/TYK2 activation^45–47^. *Shigella sonnei* inhibits all IFN signaling, including type I IFN, through effectors binding and inhibiting host cell calmodulin^48^. *L. pneumophila* specifically inhibits type I IFN signaling downstream of STAT1 and STAT2 phosphorylation via an effector secreted by its T4SS^49^.

Here we examined the role of type I IFN, specifically IFNβ, in the context of *Salmonella enterica* serovar Typhimurium (STm) infections and the capacity of STm to inhibit IFNAR-signaling in macrophage host cells. We show that IFNβ has no effect on STm growth in *ex vivo* infections of primary murine macrophages. STm infection inhibits IFNβ signaling and not IFNγ signaling in a murine macrophage reporter cell line. We show that the inhibition is at the level of IFNβ-induced STAT1 phosphorylation (pSTAT1) or further upstream in the IFNAR-signaling pathway through western blot analysis of STAT1 phosphorylation and monitoring of pSTAT1 translocation to the cell nucleus. This inhibition is independent of the T3SSs encoded by SPI-1 and SPI-2 and T6SS encoded by SPI-6. We also inhibited phagocytosis of bacteria using cytochalasin D and continued to observe reduced pSTAT1 induced by IFNAR signaling in STm-infected cells. We determined that crosstalk from TLR4 signaling mediates this inhibition as stimulation withTLR4 agonist LPS specifically inhibits IFNβ signaling, and we observed no inhibition of signaling in STm-infected TLR4-deficient BMDMs. In summary, our study reveals that although IFNβ does not affect STm growth *ex vivo*, STm infection leads to the inhibition of IFNAR-signaling in a TLR4-dependent manner.

## Results

### STm specifically inhibits type I IFN signaling

Some facultative intracellular bacterial pathogens inhibit IFNβ signaling within host cells ^47–49^. To determine if STm has this capacity, we used an *Irf3* deficient macrophage cell line (RAW264.7) that expresses Lucia luciferase, downstream of an IFN-inducible promoter (RAW-Lucia ISG-KO-IRF3). This results in a simple, luminescent readout for IFNAR and interferon gamma (IFNγ) signaling^47,49^, which signals through its own unique receptor (IFNGR)^19^. We infected RAW-Lucia ISG-KO-IRF3 cells for 30 minutes at a multiplicity of infection of 10. Subsequently, we chased the cells with 100 μg/ml gentamicin for 1 hour to kill extracellular bacteria. This was followed by 20 hours of stimulation with increasing amounts of IFNβ supplemented with 20 μg/ml gentamicin to prevent extracellular bacterial growth (Fig. 1A, left panel). Since STm infection alone induces some luminescence, we subtracted the average relative luminescent units (RLUs) produced without the addition of exogenous IFN to isolate the specific effect of STm infection on IFNAR and IFNGR signaling. While we consistently observed a reduction in RLUs in STm-infected cells regardless of IFNβ dosage used, the variability in the total RLUs between independent experiments results in a non-significant difference as determined by statistical analysis. To overcome the variability in the total RLU readouts, we normalized the results by calculating the fold change with respect to the uninfected, IFNβ-stimulated control. We then observed a statistically significant reduction in luminescence of about 30-40% (Fig. 1A, right panel). This inhibition of IFNβ signaling does not increase with multiplicity of infection (MOI) (SF1). To determine if this inhibition was specific to IFNAR, we also stimulated STm-infected cells with increasing amounts of IFNγ which signals through IFNGR (Fig. 1B). We saw a similar variability of total RLU values with IFNγ stimulation between independent experiments (Fig. 1B, left panel) but did not observe a reduction of RLUs in STm-infected cells when compared to uninfected even when the data was normalized (Fig. 1B, right panel). These data indicate that STm specifically inhibits IFNβ-induced IFNAR signaling.

**Figure 1:**
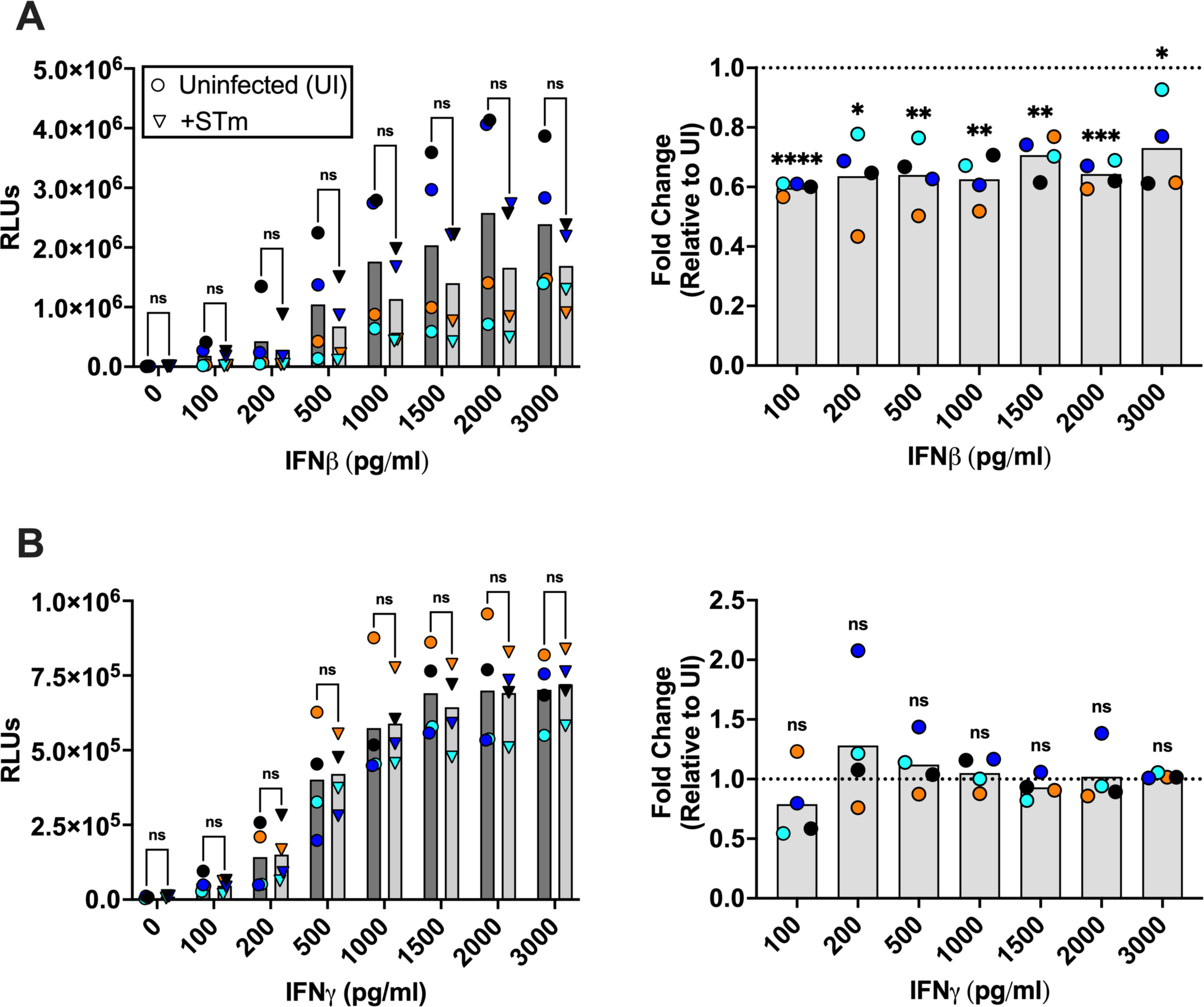
*Salmonella enterica* serovar Typhimurium (STm) infection specifically inhibits IFNβ signaling. RAW-Lucia ISG-KO-IRF3 cells were infected with STm at MOI 10 for 30 min followed by 1 hour 100 μg/ml gentamicin chase. Post-infection, cells were stimulated with IFNβ (A) or IFNγ (B) in the presence of 20 μg/ml gentamicin for 20h and the response of the gene reporter measured by luminescence and plotted as relative luminescence units (RLUs) (A and B, left panels) as well as fold change relative to the uninfected, IFNβ- or IFNγ-treated condition (A and B, right panels). Data points indicate independent biological experiments and are the average from 4 technical replicates. Statistical analysis for RLUs is multiple unpaired t-tests, and for fold change is one sample t and Wilcoxon test where each fold change mean is compared to 1. Significance is as follows: * P ≤ 0.05, ** P ≤ 0.01, *** P ≤ 0.001, **** P ≤ 0.0001.

### IFNβ has no effect on viability of STm in infected BMDMs

IFNβ has been previously shown to be protective in the context of some bacterial pathogens^31,32,50^. Its role in the context of *Salmonella* infections has been shown to be both protective^36–38^ as well as detrimental^39–44^. To determine if IFNβ is protective in the context of *ex vivo* infections, we infected BMDMs with STm for 30 minutes followed by a 1 hour gentamicin chase to kill extracellular bacteria. We then stimulated the cells with 1 ng/ml IFNβ every 12 hours in the presence of 20 μg/ml gentamicin. To account for STm infection-induced production of IFNβ, which could be acting as a confounding factor within our *ex vivo* infections, we also infected *Ifnβ^-/-^* BMDMs alongside wild type BMDMs. These infections and stimulations were followed by assessing bacterial burden through counting colony forming units (CFUs). Overall, we observed a rapid and strong restriction of STm across all conditions (Fig. 2A). While there was some variability in the initial CFUs between the conditions, we observed a 3 to 5-fold reduction in CFUs by 4 hours, and near total eradication of bacteria by 24 hours (Fig. 2A). We observed no significant difference in CFUs between experimental conditions at any timepoint. We also performed LDH release assays on the supernatants from our conditions at all timepoints and observed no significant difference in cell death across all conditions and timepoints (Fig. 2B). Thus, IFNβ does not appear to play a role in restricting STm during *ex vivo* infections of BMDMs.

**Figure 2:**
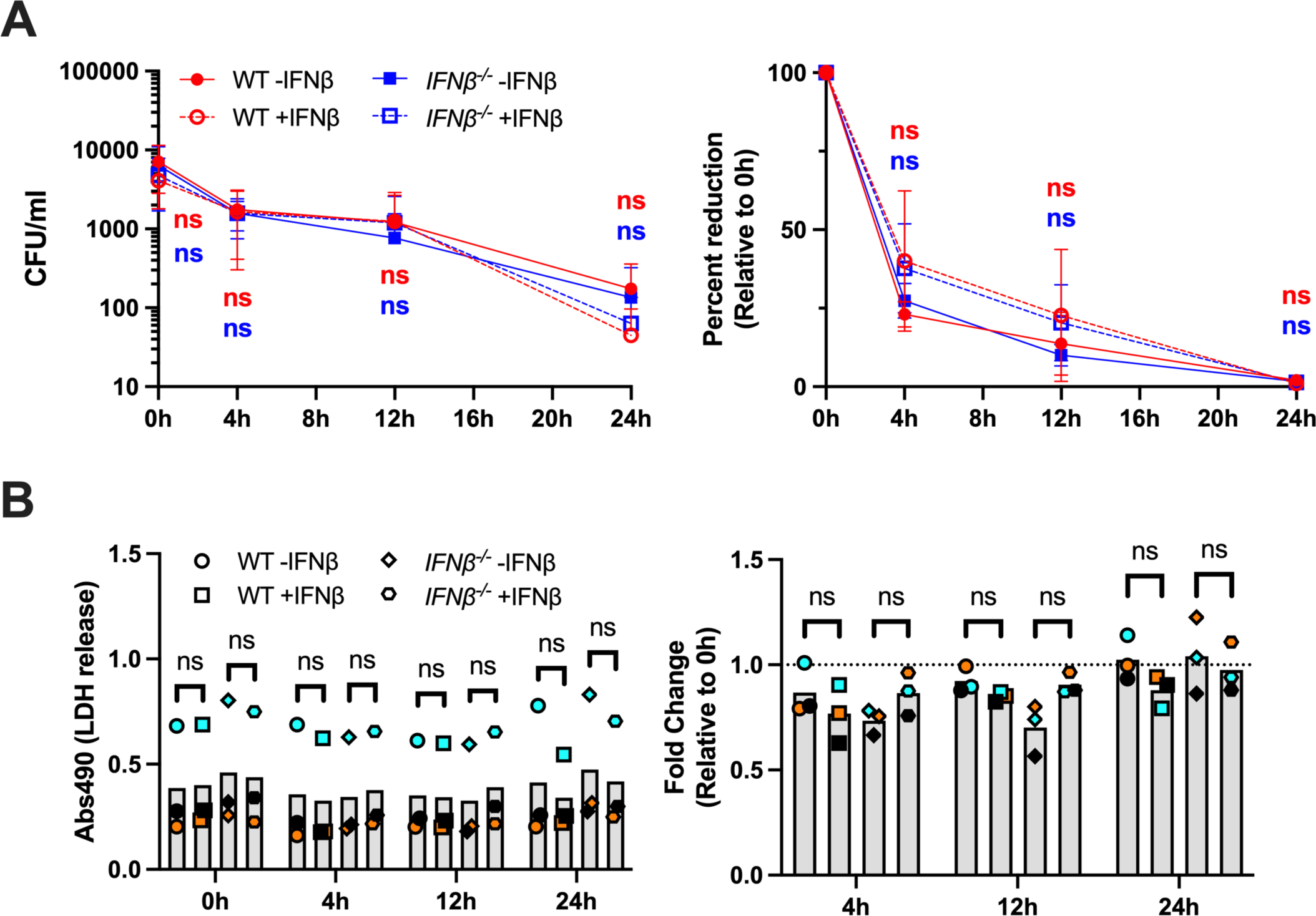
IFNβ has no effect on restriction of STm in *ex vivo* BMDM infection nor cell death induction. (A) BMDMs from wild type and *Ifnb^-/-^* mice were infected with STm at MOI 3 for 30 min min followed by 1 hour 100 μg/ml gentamicin chase. Next, cells were stimulated with 1 ng/ml IFNβ every 12h in the presence of 20 μg/ml gentamicin. Colony-forming units (CFUs) were measured at the indicated timepoints. The data shown are mean values and standard deviation of three independent infections (left panel) or the normalized to 0h post infection data (right panel). (B) LDH cell death assay was performed on cell supernatants obtained from the experiment described in panel A for each given timepoint. Data points indicate independent biological replicates (average from the 2 technical replicates) and are shown as relative absorbance at 490nm (left panel) or normalized to 0 h post infection (right panel). Statistical analysis for raw absorbance values is multiple unpaired t-test with Welch correction where each background strain stimulated with IFNβ is relative to untreated cells for a given timepoint. Statistical analysis for fold change relative to the 0h timepoint is multiple unpaired t-test with Welch correction.

### Inhibition of IFNβ signaling in STm-infected cells is upstream of STAT1 phosphorylation

The IFNβ-induced signaling pathway results in an increase in phosphorylation of STAT1 and STAT2 proteins^51^. To determine which part of the signaling pathway is inhibited by STm infection, we examined levels of phosphorylated STAT1 (pSTAT1). We infected the RAW-Lucia ISG-KO-IRF3 cells with STm at MOI of 3 for 30 minutes plus a 1 hour gentamicin chase, followed by stimulation with 200 pg/ml IFNβ for 15 minutes, 30 minutes, and 60 minutes. The cells were lysed and the levels of pSTAT1, total STAT1 and β-actin proteins were determined via western blotting (Fig. 3A). Quantification of the relative pSTAT1/β-actin and pSTAT1/STAT1 levels was performed via densitometry and normalized to the uninfected, IFNβ-treated conditions (Fig 3B). STm infection alone did not induce STAT1 phosphorylation (Fig. 3A). In STm-infected cells stimulated with IFNβ, we observed a ∼45% reduction (p = 0.024) in pSTAT1/β-actin levels at 15 minutes, ∼33% (p = 0.113) at 30 minutes, and ∼30% at 60 minutes (p = 0.158) (Fig. 3B, right panel). For pSTAT1/STAT1 levels, we measured ∼35% (p = 0.013) and ∼33% (p = 0.019) reductions in infected cells with respect to uninfected cells after 15 minutes and 30 minutes of IFNβ signaling, respectively. We observed a ∼29% reduction in pSTAT1/STAT1 levels in infected cells stimulated for 60 minutes, but it was not statistically significant (p-value = 0.058). Overall, STm-infected cells have inhibited IFNβ-induced STAT1 phosphorylation.

**Figure 3:**
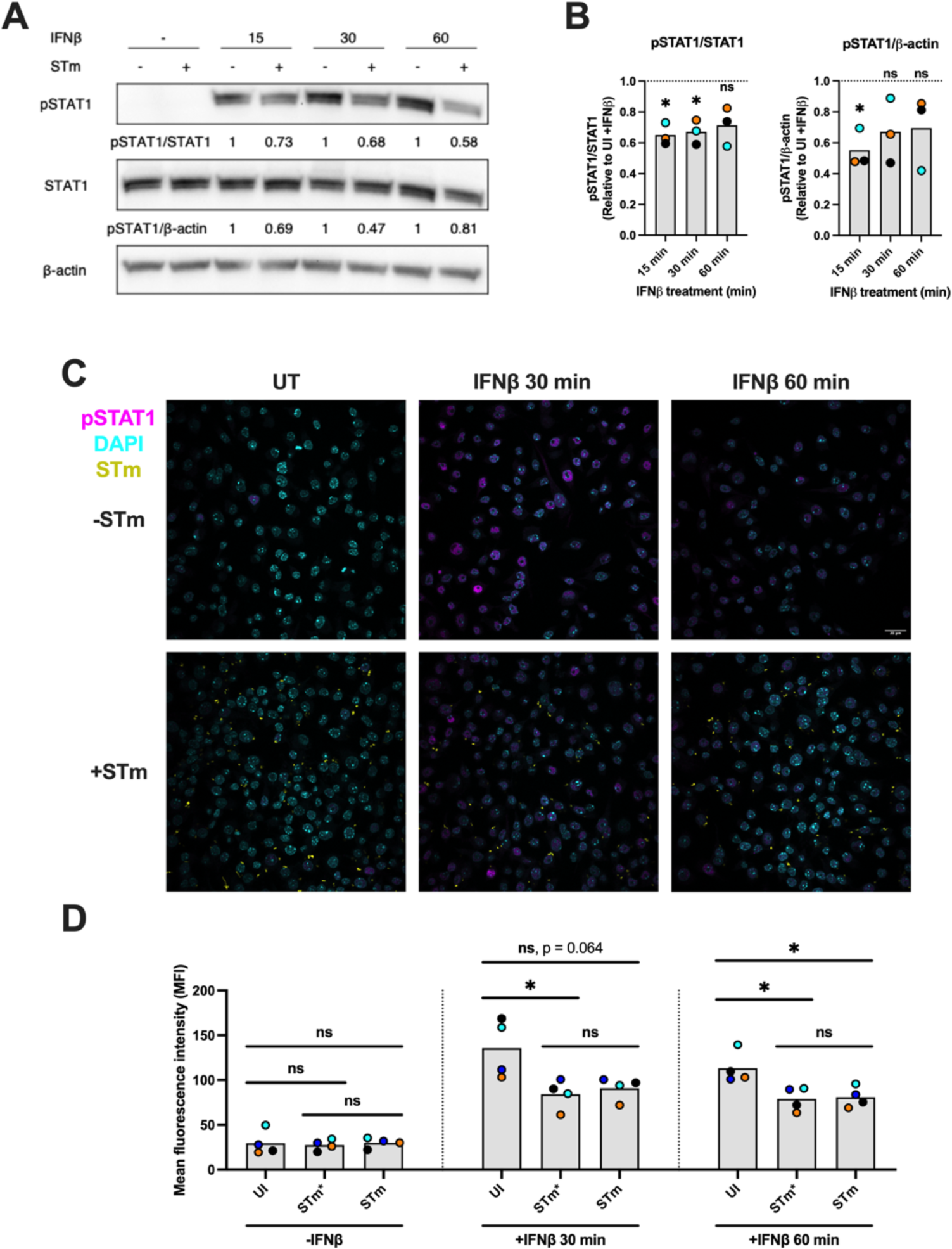
STm infection results in reduced STAT1 phosphorylation after IFNβ treatment. (A, B) RAW-Lucia ISG-KO-IRF3 cells were infected at MOI 3 for 30 min followed by 1 hour 100 μg/ml gentamicin chase. Next, cells were stimulated with 200 pg/ml IFNβ for the indicated time points in the presence of 20 μg/ml gentamicin. Whole cell lysates were collected and immunoblotted for pSTAT1, STAT1, and β-actin (A). Band densities were normalized to either STAT1 or β-actin as indicated. Densitometric ratios are shown relative to UI +IFNβ control (B). The blot shown here is representative of 3 independent infections, and individual points indicate the average from 3 independent experiments. (C) Representative images of RAW-Lucia ISG-KO-IRF3 cells infected or not with STm and treated with IFNβ or left untreated (UT). Cells were infected with yellow fluorescent protein expressing STm (STm-YFP) at MOI 3 for 30 min followed by 1 hour 100 μg/ml gentamicin chase. Following infection and chase, cells were stimulated with 1 ng/ml IFNβ for the indicated times, fixed and stained for pSTAT1 and nucleus (DAPI). (D) Mean fluorescence intensity (MFI) of pSTAT1 staining in nuclei segmented with Cellpose was quantified in uninfected (UI), infected (STm) and bystander cells (STm*). Points represent the mean of MFI values from 4 independent experiments. Statistical analysis for B is one sample t and Wilcoxon test where each fold change mean is compared to 1. Statistical analyses for D are unpaired t-test with Welch correction. Significance is indicated as such: * P ≤ 0.05, ** P ≤ 0.01.

Phosphorylated STAT1 proteins translocate to the cell nucleus to initiate transcription of interferon stimulated genes^51^. We show that STm infection inhibits STAT1 phosphorylation and in order to examine if the translocation of STAT1 proteins from the cytoplasm to the nucleus is affected, we infected RAW-Lucia ISG-KO-IRF3 with STm-YFP at MOI 3 for 30 minutes, then chased with gentamicin for 1 hour in the presence and absence of IFNβ stimulation for 30 minutes and 60 minutes followed by detection of pSTAT1 via immunofluorescence staining (Fig. 3C). We then quantified the average pSTAT1 signal within the cell nuclei using CellPose, a trainable cell segmentation algorithm, as previously reported^52^. In addition, within the infected samples, we further differentiated between infected and uninfected, bystander cells by detecting fluorescent *Salmonella* signal. Consistent with our western blot data, we observed no significant increase in average nuclear pSTAT1 with STm infection alone when compared to uninfected cells (Fig. 3D). After 30 minutes of IFNβ stimulation, we observed a 450% increase in pSTAT1 signal in uninfected cells (Fig. 3D) but only a 311% increase in STm-infected cells. This difference between uninfected and infected cells was not statistically significant (p = 0.064) by one-way ANOVA probably due to the large range of the values in the uninfected group. Interestingly, a similar reduction in pSTAT1 translocation was observed in the bystander cells (p < 0.05 compared to uninfected but p > 0.05 when compared to infected cells) (Fig. 3D). After 60 minutes of IFNβ stimulation, uninfected stimulated cells showed a 377% increase in nuclear pSTAT1, while infected cells only had a 267% increase in nuclear pSTAT1 (p < 0.05). Again, bystander cells had no significant difference in nuclear pSTAT1 with respect to their infected counterparts but were significantly reduced when compared to uninfected cells (Fig. 3D). In summary, STm infection inhibits IFNβ-induced nuclear pSTAT1 translocation regardless of bacterial uptake.

### STm inhibits type I IFN signaling independently of type III and type VI secretion systems as well as bacterial phagocytosis

STm makes extensive use of secretion systems to mediate pathogenesis and modulate the host immune response^7,53–55^. Given this, we asked if the inhibition of IFNAR signaling could be mediated by effectors from the *Salmonella* T3SSs encoded by SPI-1 or SPI-2, and the T6SS encoded by SPI-6. We infected RAW-Lucia ISG-KO-IRF3 cells with wild-type STm as well as STm mutants deficient in a specific secretion system at MOI 10 for 30 minutes. This was followed by a 1 hour gentamicin chase and stimulation with 200 pg/ml IFNβ in the presence of 20 μg/ml gentamicin. For SPI-1, deletions were made in the genes *hilA* and *hilD*, which code for transcriptional regulators of the secretion system^56–60^. For SPI-2, mutants have deletions in *ssaV* and *sseB*, essential proteins in the SPI-2 platform and translocon, respectively^61,62^. The SPI-6 mutant has a deletion in *sciG* (also called *clpV*), a chaperone required for proper effector secretion and apparatus formation^63,64^. For all these infections, we saw an inhibition of IFNAR signaling with wild type STm, approximately 30-40%. For SPI-1 mutants, we observed slightly more inhibition with respect to wild type STm, but no loss of inhibition of IFNβ signaling (Fig 4A). We also saw the same phenotype with the SPI-2 mutants (Fig. 4B) and SPI-6 mutants (Fig. 4C). In conclusion, we observed no significant difference in inhibition of IFNβ signaling between wild type STm and the different SPI mutants. We wanted to ensure that this perceived inhibition of IFNβ signaling was not due to cell death, so we performed lactose dehydrogenase (LDH) release assay at the endpoints of our infections. We observed a statistically significant increase in cell death of about 30% relative to uninfected cells in our infections (SF2A). Furthermore, we noted that this increase in cell death correlated with MOI (SF2B). Due to the significant increase in cell death observed for these infections, we decided to reduce the MOI to 3 going forward as significant inhibition was still observed at a reduced level of cell death. Additionally, we addressed the concerns of cell death contributing to this observed inhibition in our subsequent studies.

**Figure 4:**
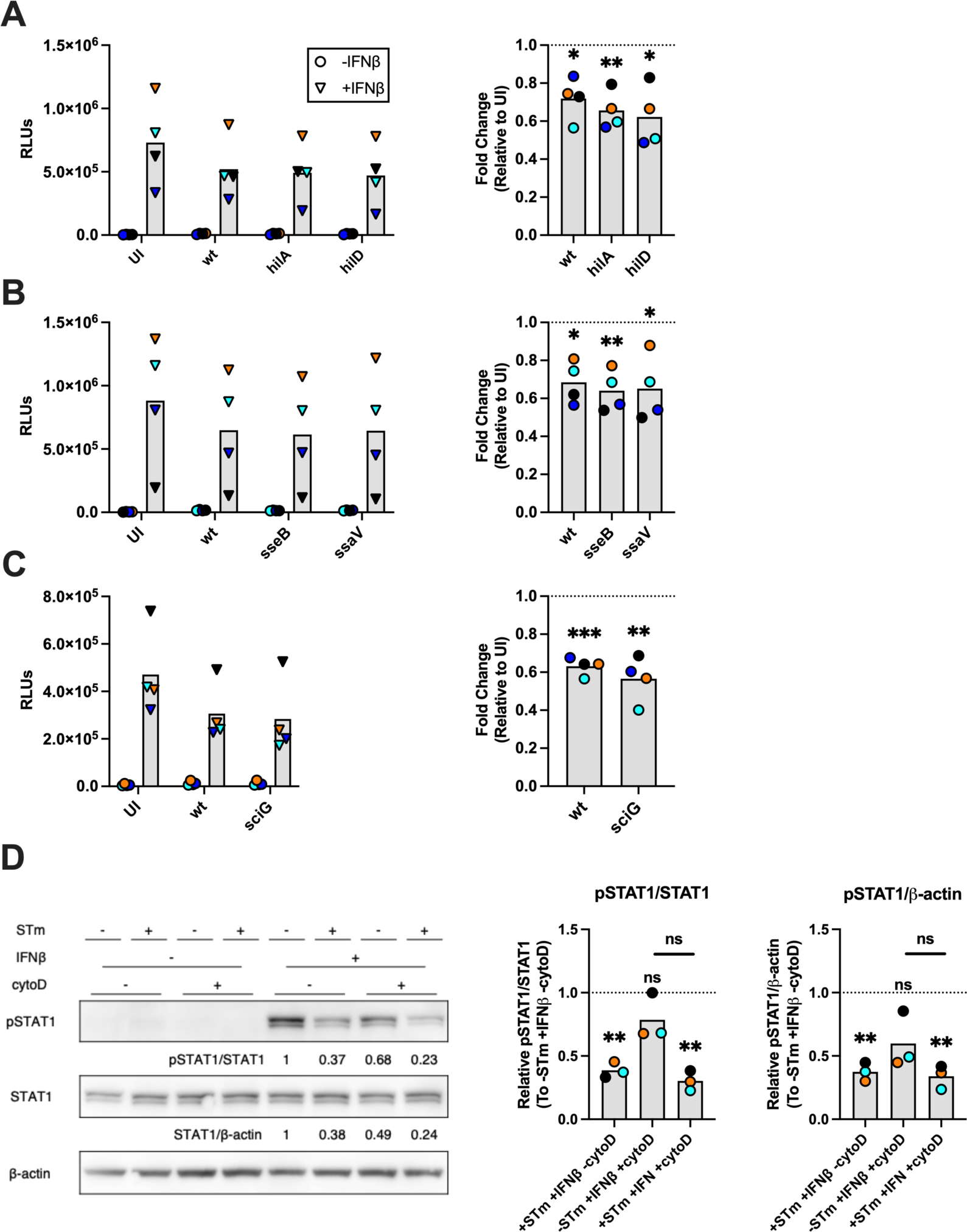
STm infection inhibits IFNβ signaling independently of type III and type VI secretion systems as well as bacterial phagocytosis. RAW-Lucia ISG-KO-IRF3 cells were infected with wild type STm or (A) SPI-1 mutants or (B) SPI-2 mutants to delete the encoded T3SS and (C) SPI-6 mutant to delete the encoded T6SS at MOI 10 for 30 min followed by 1 hour 100 μg/ml gentamicin chase. Following infection, cells were stimulated with 200 pg/ml IFNβ in the presence of 20 μg/ml gentamicin for 20h. Values are plotted as relative luminescence units (RLUs) as well as fold change relative to an uninfected IFNβ-treated control. Data points indicate independent biological experiments and are the average from 4 technical replicates. (D, E) RAW-Lucia ISG-KO-IRF3 cells were pretreated with cytochalasin D (cytoD) for 30 minutes followed by infection at MOI 3 for 30 min. CytoD was maintained on the cells at all points during the experiment. Whole cell lysates were collected and immunoblotted for pSTAT1, STAT1, and β-actin. Band densities were normalized to either STAT1 or β-actin as indicated (E). Densitometric ratios are shown relative to UI +IFNβ control (D). Statistical analysis for RLUs is multiple unpaired t-tests. For fold change and western blot quantification, statistical analyses are one sample t and Wilcoxon test where each fold change mean is compared to 1. Significance is as follows: * P ≤ 0.05, ** P ≤ 0.01, *** P ≤ 0.001.

Since bacteria-associated secretion systems are not involved in this inhibition of IFNβ signaling (Fig. 4A-C), we hypothesized that this inhibition might result from an interaction of STm with cell surface receptors. To assess if the inhibition of IFNβ signaling is indeed a cell surface signaling event, we used cytochalasin D (cytoD), an actin filament disruptor to inhibit phagocytosis within our infection model^65–67^. We pretreated the cells with cytoD at 10 μM for 30 minutes followed by STm infection at MOI 3 for 30 minutes with cytoD maintained on the cells at all points of the infection. We confirmed that cytoD inhibits phagocytosis in the cells using flow cytometry following infection with a fluorescent strain of STm constitutively expressing yellow fluorescent protein (STm-YFP). We observed a two-fold increase in mean fluorescence intensity (MFI) in the untreated, infected experimental group when compared to uninfected cells (SF3). This STm infection-mediated increase was reversed to the baseline uninfected MFI when cells were treated with cytoD. This indicated that phagocytosis of STm was disrupted by treatment with cytoD.

To determine if the inhibition of IFNβ signaling is phagocytosis-independent, we pretreated RAW-Lucia ISG-KO-IRF3 cells with cytoD followed by STm infection at MOI 3 for 30 minutes plus 1 hour gentamicin chase, then treatment with IFNβ (200 pg/ml) for 30 minutes. CytoD was kept on the cells at all points of the experiment. The cells were then lysed and pSTAT1, STAT1, and β-actin levels were examined via western blot (Fig. 4D). Like our previous results, we measured no induction of pSTAT1 levels in the RAW-Lucia ISG-KO-IRF3 cells upon STm infection alone or plus cytoD treatment (Fig. 4D). When stimulated with IFNβ, we saw an increase in pSTAT1/STAT1 and pSTAT1/β-actin levels, which was reduced with STm infection by about 37% and 39% respectively. When stimulated with cytoD and IFNβ, we saw a reduction in pSTAT1/STAT1 and pSTAT1/β-actin with respect to the uninfected, IFNβ-stimulated control. For pSTAT1/STAT1, this reduction was about 21%. For pSTAT1/β-actin, this reduction was about 40%. When cytoD treatment and IFNβ stimulation were combined with STm infection, we continued to see an inhibition of pSTAT1/STAT1 and pSTAT1/β-actin, with reductions of about 70% for pSTAT1/STAT1 and about 66% for pSTAT1/β-actin. We did not expect reduced pSTAT1/STAT1 and pSTAT1/β-actin with cytoD treatment alone, but this may not surprising given previous research implicating endocytosis of IFNAR as an important part of signaling^68,69^, and actin disruption via cytoD could be affecting this. We also examined if this inhibition was dependent on existing factors within STm by heat inactivating STm cultures at 42 °C and 65 °C for 30 minutes followed by our typical infection protocol (SF4). We continued to observe inhibition in infections involving the heat inactivated STm and there was no significant difference in inhibition when compared to the non-heat inactivated STm. Overall, inhibition of IFNβ signaling in STm infection is independent of bacterial phagocytosis as well as the activity of STm type III and type VI secretion systems.

### LPS stimulation specifically inhibits IFNβ signaling

With bacterial phagocytosis and uptake ruled out as being important for STm-mediated IFNAR-signaling inhibition, we hypothesized that the inhibition is mediated by host innate immune signaling, specifically TLR4 signaling. Indeed, previous research indicates that TLR7/8 signaling can inhibit IFNα signaling^70^. To test this hypothesis, we pretreated RAW-Lucia ISG-KO-IRF3 cells with LPS, the ligand for TLR4, at 50 ng/ml for 4 hours followed by 200 pg/ml IFNβ for 20 hours (without the removal of LPS). We then collected supernatants to measure luminescence and cell death. Similar to STm infection (Fig. 1), LPS induced some luminescence independently of any exogenous IFNβ (Fig. 5A). To account for this, we subtracted the average RLUs produced with LPS alone to isolate the specific effect it has on IFNAR signaling followed by determining the fold change. We observed a consistent reduction in raw RLUs as well as fold change, with an average reduction of about 50% with respect to the -LPS +IFNβ control. To determine if this inhibition extends to other IFN signaling, we also stimulated the cells with 500 pg/ml IFNγ for 20 hours following a 4 hour stimulation with 50 ng/ml LPS. In contrast to IFNAR signaling, we observed an increased induction of raw RLUs as well as an approximate 1.8 fold increase with respect to -LPS +IFNγ control (Fig. 5B). For both IFNAR and IFNGR signaling, we saw no significant increase in cell death (Fig. 5C+D). In summary, LPS stimulation inhibits IFNβ signaling but not IFNγ signaling.

**Figure 5:**
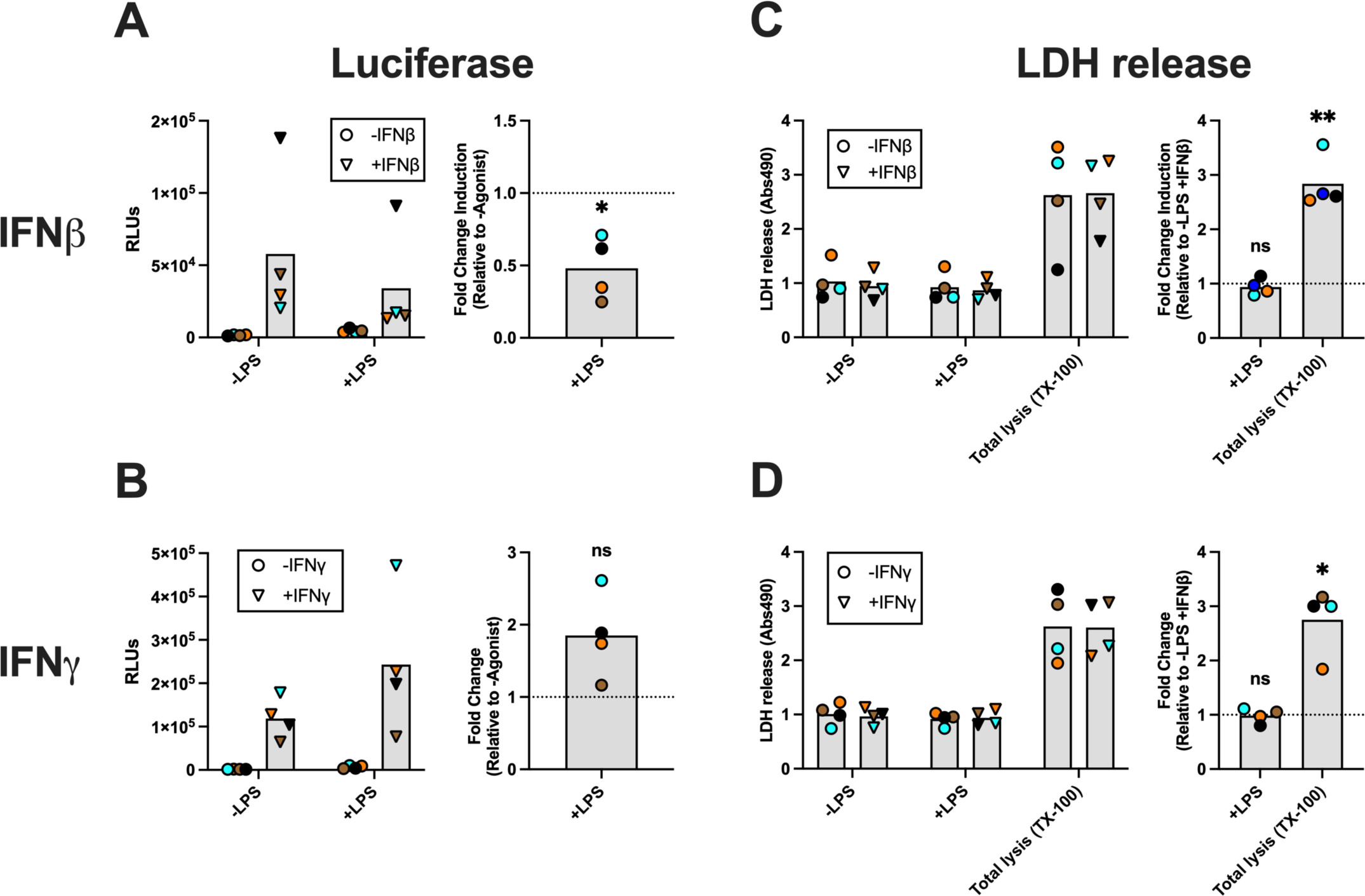
LPS stimulation specifically inhibits IFNβ signaling without affecting host cell death. RAW-Lucia cells were pre-treated with 50 ng/ml LPS for 4 hours followed by an additional 20 hours along with 200 pg/ml IFNβ (A) or 500 pg/ml IFNγ (B). Values are plotted as relative luminescence units (RLUs) as well as fold change relative to their respective -LPS +IFN control. (C+D) Supernatants were collected and analyzed for endpoint cell death from the IFNβ and IFNγ experiments (A+B) using LDH release assay. Values are plotted as absorbance at 490 nm (Abs490) as well as fold change relative to their respective -LPS +IFN control. Individual points indicate independent biological experiments and are the average from 4 technical replicates. Statistical analysis for each panel is one sample t and Wilcoxon test where each fold change mean is compared to 1. Statistical significance is as follows: * P ≤ 0.05, ** P ≤ 0.01, *** P ≤ 0.001.

### STm infection inhibits IFNβ signaling in a TLR4-dependent manner

To determine if the kintetic of the LPS-mediated inhibition, we stimulated RAW-Lucia ISG-KO-IRF3 cells with 50 ng/ml LPS from time points ranging from 15 minutes to 8 hours, after which 200 pg/ml IFNβ was added to the cells for 30 minutes. We then lysed the cells and examined the levels of pSTAT1, STAT1, and β-actin via western blot (Fig. 6C). The results of three independent experiments were quantified and we observed significant reductions in pSTAT1/STAT1 levels with concurrent LPS/FNβ stimulation for most time points, with the inhibition being most robust at 30 minutes and 1 hour post-LPS stimulation (Fig. 6A+B). We saw similar results with pSTAT1/β-actin ratios, but the inhibition is not statistically significant at 4 hours and 8 hours post-LPS stimulation (Fig. 6B). We also examined if LPS could inhibit IFNγ-induced STAT1 phosphorylation by stimulating RAW-Lucia ISG-KO-IRF3 cells with 50 ng/ml LPS for 30 minutes and 1 hour followed by stimulation with 500 pg/ml IFNγ for 30 minutes (Fig. 6C). This was followed by lysis and western blotting for of levels of pSTAT1, STAT1/ and β-actin. Over three independent experiments, we observed no significant reduction of IFNγ-induced pSTAT1/STAT1 levels or pSTAT1/β-actin with LPS stimulation (Fig. 6C+D). This supports our earlier observations that STm infection (Fig. 1) and LPS stimulation alone (Fig. 5) specifically inhibits IFNβ-induced signaling upstream of STAT1 phosphorylation.

**Figure 6:**
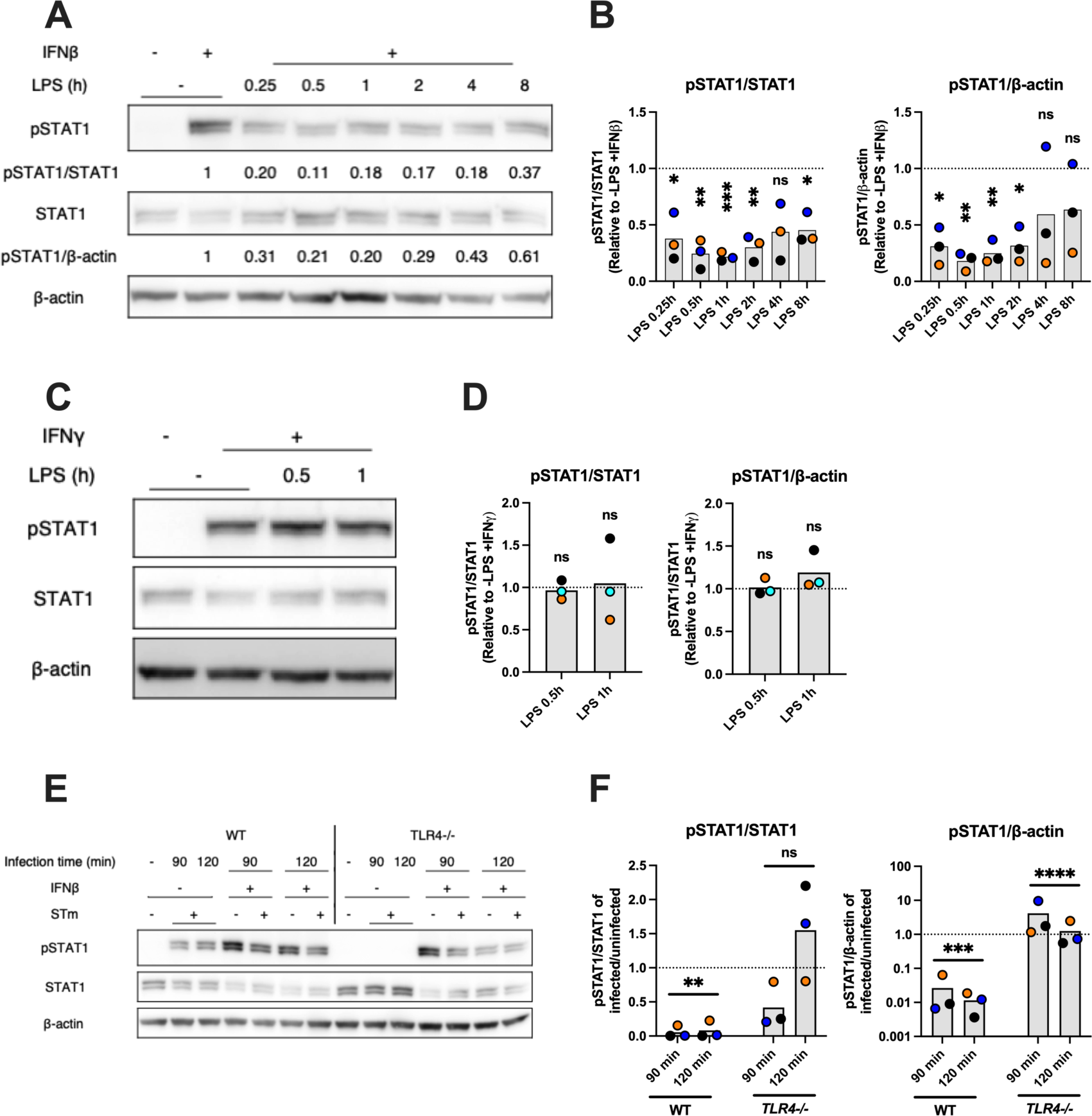
STm infection inhibits IFNβ signaling in a TLR4-dependent manner. (A-D) RAW-Lucia ISG-KO-IRF3 cells were pretreated with 50 ng/ml LPS for the indicated time points followed by stimulation with either (A, B) 200 pg/ml IFNβ or (C, D) 500 pg/ml IFNγ for 30 minutes. Whole cell lysates were collected and immunoblotted for pSTAT1, STAT1, and β-actin. Band densities were normalized to either STAT1 or β-actin as indicated. Densitometric ratios are shown relative to respective -LPS +IFNβ or -LPS +IFNγ control. Each blot is representative of 3 independent infections, and individual data points (B, D) indicate independent experiments. (E, F) BMDMs were infected with STm at MOI 3 for 30 min followed by 1 hour 100 μg/ml gentamicin chase. Cells were incubated for an additional 30 min or 60 min followed by 200 pg/ml IFNβ stimulation for 30 min (all in the presence of 20 μg/ml gentamicin). This results in 90 minutes and 120 minutes post-infection time points. Whole cell lysates were collected and immunoblotted for pSTAT1, STAT1, and β-actin. Band densities were normalized to either STAT1 or β-actin as indicated by equation in methods section. Blot is representative of 3 independent infections, and individual data points (F) indicate independent experiments. Statistical analyses are one sample t and Wilcoxon test where each ratio is compared to 1. Statistical significance is as follows: ** P ≤ 0.01, *** P ≤ 0.001, **** P ≤ 0.0001.

Next, we infected BMDMs from *Tlr4^-/-^* mice in parallel with BMDMs from wild type (C57BL/6) mice with STm at MOI3 to determine if TLR-4 signaling is important for inhibition in the context of an infection. We infected the cells for 30 minutes followed by a 1 hour gentamicin chase and since we wanted to determine if this inhibition occurred at prolonged initial timepoints, so the infected cells were rested for an additional 30 minutes or 60 minutes (with 20 μg/ml gentamicin) totaling in 90 minutes or 120 minutes of infection. This was then followed by then stimulation with 200 pg/ml by IFNβ stimulation for 30 minutes. Cells were lysed and levels of pSTAT1, STAT1, and β-actin were examined via western blot (Fig. 6E). Unlike the RAW-Lucia ISG-KO-IRF3 cells, we observed increased pSTAT1 in STm-infected wild type BMDMs but not in the infected *Tlr4^-/-^* BMDMs (Fig. 6E). Due to these differences between the wild-type and *Tlr4^-/-^* BMDMs in induction of pSTAT1 after STm infection, we adjusted our quantification method to account for that. We first calculated the ratio of pSTAT1 signal with IFNβ stimulation between infected cells and uninfected cells, then divided this value by the ratio of STAT1 or β-actin signal with IFNβ stimulation between infected cells and uninfected cells. The resulting pSTAT1/STAT1 and pSTAT1/β-actin values were then used for comparison. If the value is less than 1, that indicates signaling inhibition occurred. If the value is greater than 1, that indicates signaling induction occurred. In wild type BMDMs, we observed pSTAT1/STAT1 mean values of 0.055 and 0.081 at 90 minutes and 120 minutes post-chase, respectively (Fig. 6E+F). Both values indicate statistically significant inhibition of IFNβ-induced signaling. In *Tlr4^-/-^* BMDMs, we observed pSTAT1/STAT1 mean values of 0.42 at 90 minutes post-chase and 1.56 at 120 minutes post-chase, respectively (Fig. 6E+F). While the mean pSTAT1/STAT1 value at 90 minutes post-chase is less than 1, there is a high degree of variability that results in it being not statistically significant. The results at 120 minutes post-chase also exhibit high variability, which may not be surprising given the knockout of such an important and broadly activating innate immune signaling pathway. Indeed, knocking out TLR4 already alters STAT1 phosphorylation with STm infection (Fig. 6E)

We also observed a statistically significant reduction in pSTAT1/β-actin levels in wild type BMDMs with STm infection in the context of IFNβ-induced IFNAR signaling. We observed pSTAT1/β-actin values of 0.026 and 0.011 at 90 minutes and 120 minutes post-chase, respectively (Fig. 6E+F). In the *Tlr4^-/-^* BMDMs, we observed pSTAT1/β-actin values of 4.15 at 90 minutes post-chase and 1.25 at 120 minutes post-chase (Fig. 6E+F). Stimulating wild type BMDMs with LPS alone for 1 hour followed by a 15 min, 30 min or 60 min IFNβ stimulation resulted in reduced pSTAT1 levels compared to the non IFNβ-treated cells (SF5). Interestingly, LPS stimulation alone did not result in increased pSTAT1 in contrast to STm-infected BMDMs (SF5). This indicates that TLR4 signaling is necessary but not sufficient for STAT1 phosphorylation. We thus demonstrate in two independent experimental approaches that LPS signaling can inhibit subsequent IFNβ-mediated signaling which is consistent with STm-mediated inhibition of IFNβ-mediated signaling being dependent on TLR4.

## Discussion

While Type I IFNs, like IFNβ, generally have a beneficial role in viral infections^25,26,29^, their role in bacterial infections can vary, exhibiting either advantageous or detrimental effects. This is dependent on factors such as the bacterial pathogen and site of infection^31–33^. In the context of STm infection, the role of type I IFN is not clearly understood, with conflicting results indicating either a detrimental^39–41,43,44^ or a protective^38,42^ role. Here, we show that IFNβ has no effect on the growth of STm in *ex vivo* macrophage infections (Fig. 2). We demonstrate a strong restriction of STm growth in BMDMs regardless of the presence of endogenous or exogenous IFNβ. A limitation of this result is that the absence of STm growth in BMDMs derived from C57BL/6 mice may hinder the detection of an enhanced cell-autonomous defense following the addition of IFNβ. In future studies, BMDMs obtained from a mouse strain that supports intracellular STm growth could reveal a protective role of IFNβ. Nevertheless, our results are consistent with previous research indicating that the priming of murine macrophages with IFNβ prior to STm infection had no effect on CFU recovery post-infection^40^. IFNβ priming did downregulate neutrophil chemokine and proinflammatory cytokine secretion, hinting at a possible role during *in vivo* infections when multiple host cell types are involved in the immune response^40^. IFNβ is protective in murine gastrointestinal infection by STm^37^. In that model, STm-infected intestinal macrophages produce IFNβ, which restricts STm growth and dissemination by activating bystander natural killer cells to produce IFNγ. IFNγ then exerts antimicrobial effects and stimulates phagocytes to release CXC chemokines, attracting other immune cells^37^. Conversely, in a murine systemic STm infection model (via intravenous tail vein injection), type I IFN proves detrimental since *Ifnar^-/-^* are more resistant to STm infection^39,42^. The mechanism of resistance involves a decrease in necroptosis of *Ifnar^-/-^* macrophages after STm infection^39^ as their macrophages resist necroptotic cell death induced by type I IFN signaling. The current data suggest that type I IFN may play a protective role in the context of *in vivo* gastrointestinal infections via the oral route but can be detrimental in the context of a systemic infection. Complicating matters is a report that IFNβ production in mice infected with influenza virus increases susceptibility to STm-mediated gastroenteritis following oral infection^41^. Consequently, further studies are needed to fully elucidate the role of IFNβ during in vivo STm infections.

Given the beneficial role of IFNβ in viral infections, inhibition of IFNβ signaling by viruses is common^28,71–73^. In comparison to viruses, there are few examples of bacterial pathogens that can inhibit IFNβ signaling. *Shigella sonnei* inhibits all types of IFN signaling through its secreted effector, OspC, which acts by binding to and inhibiting host cell calmodulin. Consequently, calmodulin kinase II activation is inhibited which leads to reduced STAT1 phosphorylation^48^. In addition, *Mycobacterium tuberculosis* and *Legionella pneumophil*a infection specifically inhibits IFNβ signaling via an unknown mechanism^47,49^. Here, we show that STm infection inhibits IFNβ-induced IFNAR signaling. Interestingly, unlike *S. sonnei* that inhibit IFNβ signaling through secreted effectors^48,49^, we show that STm infection inhibits IFNβ signaling through crosstalk with TLR4 signaling. This pathway may be activated by other Gram-negative bacteria that trigger TLR4 signaling. If TLR2 signaling can also inhibit IFNAR signaling, it opens the possibility of involvement by Gram-positive bacteria as well. This is not the first time TLR signaling has been implicated in inhibitory crosstalk with type I IFN signaling^70^. However, to our best knowledge, this is the first time that TLR signaling was shown to inhibit IFNβ signaling during bacterial infection of macrophages.

Negative regulation of type I IFN signaling is complex, with regulators exerting their functions at several steps in the signaling pathway^15,24,22^. Some negative regulators such as the suppressor of cytokine signaling (SOCS) family, specifically SOCS1, as well as ubiquitin-specific peptidase 18 (USP18), and protein inhibitor of activated STAT (PIAS) are not constitutively expressed and thus inhibit IFNβ signaling at later timepoints^19–21^. In contrast, protein tyrosine phosphatases are regulators of type I IFN signaling that can act immediately and can have positive or negative regulatory functions^15,24^. The earliest timepoint we analyzed during STm infection and observed inhibited IFNβ signaling is about 90 min after the macrophages first encountered bacteria (Fig. 3A+B). This might be enough time for a regulator such as USP18 to be available for inhibition, since, for example, the β2-integrin-mediated inhibition of IFNAR-signaling was apparent after only one hour of stimulation with the β2-integrin ligand fibrinogen^74^. Nevertheless, our data showing that LPS inhibits signaling after only 15 min suggests that other factors are involved. Another argument against SOCS1 and USP18 is that they do not act specifically on IFNAR signaling^22^, which is what we observed for STm-mediated IFNβ signaling inhibition (Fig. 1). A likely candidate is the SH2-containing tyrosine phosphatase 2 (SHP-2) which inhibits IFNAR signaling after of only 15 min of activation via TLR7 and TLR8 signaling^70^.

In conclusion, we show that STm infection inhibits IFNβ signaling via crosstalk with the TLR4 signaling pathway and that LPS alone can stimulate TLR4 to inhibit IFNβ signaling. Multiple signaling pathways crosstalk with IFNAR signaling which could be how cells “fine-tune” their immune response, leading to improved or worsened immune responses depending on the pathogen and the host^75^. Many questions remain, such as the molecular mechanism of TLR4 signaling that leads to IFNβ signaling inhibition, the relevance of this inhibition during *in vivo* infections with STm, and if other bacteria engage the same inhibitory pathway.

## Methods

### Cell culture and mice

The RAW-Lucia ISG-KO-IRF3 cells (Invivogen) were grown in Dulbecco’s modified Eagle medium (DMEM, Gibco or ATCC, 11965092 or 30-2002 respectively) with 10% heat inactivated fetal bovine serum (FBS, Gibco, A52567-01).

*Tlr4^-/-^* mice^76^ and *Ifnβ^-/-^* mice were provided by Dr. Stefanie Vogel. The *Ifnβ^-/-^* mice were originally generated by Dr. E Fish (University of Toronto) and have been previously described^77^. C57BL/6J mice were either provided by Dr. Stefanie Vogel or obtained from The Jackson Laboratory. Animal studies were approved by Institutional Animal Care and Use Committee and conducted in accordance with the NIH. BMDMs were prepared as previously described^49^. In brief, bone marrow cells were flushed from mouse femurs and tibias followed by culturing in supplemented DMEM for 6 days prior to infection.

### Bacteria

*Salmonella enterica* serovar Typhimurium strain 12408s and *ΔssaV* and *ΔsseB* (SPI-2 deficient) mutants were provided by Dr. Sophie Helaine^78^. The *ΔhilA* and *ΔhilD* (SPI-1 deficient), *ΔsciG* (SPI-6 deficient, also goes by *ΔclpV*), and yellow fluorescent protein-expressing *Salmonella* (STm-YFP) were provided by Dr. Jiqiang Ling^52,60,79^. The SPI-1 mutants and SPI-6 mutant were constructed as previously described^80^ using chloramphenicol (Chl) as the resistance marker. Briefly, the Flp recombination target (FRT)-flanked Chl cassette was amplified by PCR from plasmid pKD3. The resulting PCR product was integrated into STm chromosome by lambda-red recombination. The positive clones were selected on Luria-Burtani (LB) plates containing 25 μg/ml chloramphenicol at 30°C and verified by PCR. Primer sequences used for mutagenesis are listed in Table S1. Bacterial strains were grown in LB medium supplemented with chloramphenicol (25 ug/ml) for the SPI-1 and SPI-6 mutants, or kanamycin (50 ug/ml) for the SPI-2 mutants respectively. *Salmonella* cultures were grown overnight at 37 °C shaking at 225 rpm or 250 rpm followed by resuspension in phosphate-buffered saline and opsonization in a suspension of DMEM with non-heat inactivated fetal bovine serum and normal mouse serum (Jackson ImmunoResearch, 015-000-120) for 20 minutes.

### Reporter cell assays

RAW-Lucia ISG-KO-IRF3 cells were seeded at 1 x 10^5^ cells per well in quadruplicate in 96-well tissue culture-treated plates overnight. The medium used was DMEM without serum to reduce background activation of the ISRE-luciferase gene cassette. Cells were infected at an MOI of 10 for 30 minutes at 37 °C after centrifugation at 100 x g for 5 minutes. Cells were washed three times with PBS followed by 1 hour incubation with complete DMEM containing 100 μg/ml gentamicin (Corning, 30-005-CR). This medium was aspirated and replaced with complete medium containing 20 μg/ml gentamicin and IFNβ (R&D Systems, 8234-MB-010) as indicated for 20 hours. For LPS (Invivogen, tlrl-eblps) stimulations, cells were stimulated at a concentration of 50 ng/ml for the indicated times in DMEM with heat inactivated fetal serum followed by the addition of IFNβ on top the existing LPS (no aspiration) also for the indicated times.

### Western blot analysis

RAW-Lucia cells were seeded at 1 x 10^6^ cells per well in triplicate in 24-well tissue culture-treated plates overnight. The medium used was complete DMEM. For infections, cells were in infected at an MOI of 3 and infected as previously described above. For cytochalasin D infections, cells were first pretreated with 10 μM cytochalasin D for 30 minutes followed by our usual infection protocol, but with 10 μM cytochalasin D maintained on the cells for all points of the experiment. Cells were rinsed with PBS twice prior to lysis. Whole cell lysates were made by lysing cells with RIPA buffer containing Halt protease and phosphatase inhibitor cocktail (ThermoFisher, 78440) and Dnase I (ThermoFisher, 90083). Protein concentrations were determined with Pierce BCA Protein Assay Kit (ThermoFisher, 23225) followed by SDS-PAGE and immunoblotting. Antibodies were detected using SuperSignal West Pico chemiluminescent substrate (ThermoFisher, 34580) or SuperSignal West Femto chemiluminescent substrate (ThermoFisher, 34096).

The equation used for quantification of pSTAT1/STAT1 and pSTAT1/β-actin signal is as follows (UI = uninfected):

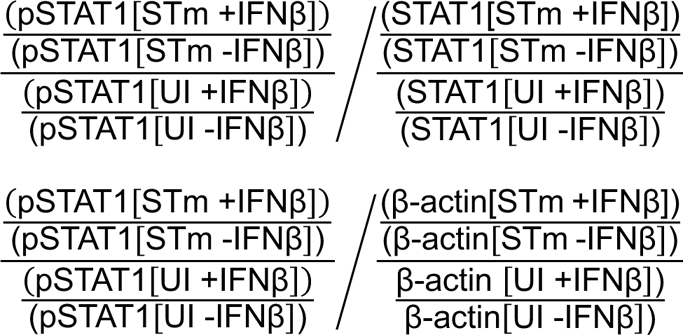

### Bacterial survival

BMDMs were seeded at 5 x 10^5^ cells per well in duplicate in 24-well tissue culture-treated plates overnight. The medium used was DMEM with 10% heat-inactivated FBS and 20% L929-conditioned medium (BMDM growth medium). Cells were infected at an MOI of 3 as described above. IFNβ was added at 1 ng/ml every 12 hours. At the indicated timepoints, cells were washed with PBS three times followed by addition of 0.1% Triton X-100 in PBS for 5 minutes. The duplicate wells were then pooled, serially diluted, plated in duplicate onto LB plates, and incubated at 37 °C for approximately 16 hours. Bacteria were enumerated based on colony forming units (CFUs).

### Immunofluorescence

RAW-Lucia cells were seeded at 1 x 10^6^ cells onto 12 mm diameter #1 coverslips (Electron Microscopy Sciences, 72231-01) in duplicate in complete DMEM overnight. Following infection with STm-YFP and IFNβ stimulation, cells were rinsed three times with PBS followed by fixation in 4% paraformaldehyde at room temperature for 30 minutes. Paraformaldehyde was aspirated and PBS was added to the cells for 5 minutes for a total of three washes. To permeabilize the cells, −20 °C methanol was added to the cells and incubated at −20 °C for 20 minutes. The methanol was removed and PBS containing 10% normal goat serum (Jackson ImmunoResearch, 005-000-121) and 0.3% Triton X-100 was added and incubated at room temperature for 1 hour. This suspension was removed and replaced with PBS containing 0.3% Triton X-100, 0.1% bovine serum albumin (P/B/T), and the pSTAT1 antibody. This suspension was left on the cells overnight at 4 °C.

Following primary antibody incubation, cells were incubated with PBS three times as previously described in this section followed by incubation with P/B/T containing goat anti-rabbit antibody conjugated to Alexa Fluor 647 (Jackson ImmunoResearch, 111-605-144) for 2 hours at room temperature protected from light. Cells were incubated and washed thrice with PBS as previously described followed by incubation with PBS containing DAPI (NucBlue, Invitrogen, R37606) for 30 minutes. Cells were incubated and washed thrice with PBS followed by mounting with Prolong Diamond Antifade Mountant (Invitrogen, P36961) for approximately 18-24 hours before imaging.

### Image analysis

The bacterial burden quantification, infection rate, and total fluorescence intensity of nuclear pSTAT1 signal was performed in high throughput using PyimageJ^81^ running in JupyterLab 4.0. The code used for this study is available on GitHub (https://github.com/jaugenst/nucleus_fluorescence_quantification) and provided as supplementary material. The nuclei segmentation model trained in Cellpose^82^ is also available on GitHub. The method was derived and extended from the BBQ method published previously^52^. Briefly, after the training of a custom model (also available on GitHub), cell nuclei were detected using Cellpose. Nuclear ROIs were used to perform Voronoi segmentation and to collect the total fluorescence intensity of bacteria per cell defined by Voronoi area. This yielded the relative bacterial burden of the cells. The nuclei ROIs were applied to the pSTAT1 channel, and mean fluorescence intensity (MFI) within those ROIs was measured.

### Cell death assays

Toxilight Bioassay Kit (Lonza, LT17-127) or LDH-Glo Cytotoxicity Assay (Promega, J2380) were used to determine cell death. Toxilight Bioassay Kit measures adenylate kinase (AK) in cell supernatant due to membrane damage. CytoTox 96 Non-Radioactive Cytotoxicity Assay measures lactose dehydrogenase (LDH) in the cell supernatant due to membrane damage. Both assays were performed per their respective manufacturer protocol.

### Statistical analysis

All statistical analyses were done using GraphPad Prism 10 with the indicated tests. Significance is indicated by a p-value less than or equal to 0.05.

## Acknowledgments

Purchase of the Zeiss LSM 980 Airyscan 2 was supported by Award Number 1S10OD025223-01A1 from the National Institute of Health. We sincerely thank the Imaging Core Facility in the department of Cell Biology and Molecular Genetics at the University of Maryland, College Park, and Amy Beaven for assistance in using this microscope. Additionally, we are grateful for Dr. Stefanie Vogel, Dr. Katharina Richard, and Dr. James Barrett for their contributions of mice for this study. We would like to acknowledge the use of ChatGPT, a language model developed by OpenAI, for assistance in revising and refining sentence syntax in this manuscript.

